# Origins and geographic diversification of African rice (*Oryza glaberrima*)

**DOI:** 10.1101/398693

**Authors:** Margaretha A. Veltman, Jonathan M. Flowers, Tinde R. van Andel, M. Eric Schranz

## Abstract

Rice is a staple food for the majority of our world’s growing population. Whereas Asian rice (*Oryza sativa* L.) has been extensively studied, the exact origins of African rice (*Oryza glaberrima* Steud.) are still contested. Previous studies have supported either a centric or a non-centric origin of African rice domestication. Here we review the evidence for both scenarios through a critical reassessment of 206 whole genome sequences of domesticated and wild African rice. While genetic diversity analyses support a severe bottleneck caused by domestication, signatures of recent and strong positive selection do not unequivocally point to candidate domestication genes, suggesting that domestication proceeded differently than in Asian rice – either by selection on different alleles, or different modes of selection. Population structure analysis revealed five genetic clusters localising to different geographic regions. Isolation by distance was identified in the coastal populations, which could account for parallel adaptation in geographically separated demes. Although genome-wide phylogenetic relationships support an origin in the eastern cultivation range followed by diversification along the Atlantic coast, further analysis of domestication genes shows distinct haplotypes in the southwest - suggesting that at least one of several key domestication traits might have originated there. These findings shed new light on an old controversy concerning plant domestication in Africa by highlighting the divergent roots of African rice cultivation, including a separate centre of domestication activity in the Guinea Highlands. We thus suggest that the commonly accepted centric origin of African rice must be reconsidered in favour of a non-centric or polycentric view.

## Introduction

### History and relevance

Rice is the world’s most important cereal crop. As a staple food for more than half of the world’s seven billion people, it is of crucial importance in providing food security for an exponentially growing population. Like other cereals, rice has been domesticated by humans multiple times independently. Yet, unlike other cereal species, these domestication events have origins on different continents – one in Asia, and one in Africa. Although the exact history of rice in Asia is still disputed, it is clear that Asian rice (*Oryza sativa* L.) was domesticated from a single wild species (*Oryza rufipogon* Griff.) approximately 9000 years ago [1]. In contrast, African farmers domesticated rice from another progenitor, *Oryza barthii* A. Chev., approximately 3000 years ago. This event resulted in a species that is now recognised as *Oryza glaberrima* Steud. [2].

Asian and African rice have distinct phenotypic characteristics: their grains differ in colour, size, shape and taste. Whereas Asian rice can be milled mechanically, facilitating large-scale production, African rice grains break easily and have to be milled manually with a mortar and pestle. These characteristics have favoured the cultivation of Asian rice over African rice in large parts of the world. In Africa, *O. glaberrima* has largely been replaced by Asian rice, even though African rice is more resistant to abiotic stresses and is often preferred for its taste and its diversity in maturation time [3]. In addition, African rice continues to survive in a ritual context, used in ritual offerings to honour the ancestors, rather than for consumption [4].

Globalisation places these local cultural traditions and the neglected species associated with them under threat [5]. In addition, food demand is rising in many African countries as a result of the growing population, a trend which is reflected in annual rice consumption [6]. Yet, food security is increasingly under pressure from ongoing land use and climate change, limiting the availability of suitable crop land [7]. Both processes have accelerated the shift in cultivation from local African varieties to the more productive Asian varieties. As a result, many traditional landraces of *O. glaberrima* are disappearing, or have already disappeared [3].

Even though Asian rice has higher yields, the diminishing genetic diversity of African rice may lead to the loss of other important agronomic traits (such as salt tolerance or blight resistance) that are not represented in *O. sativa* [3]. Loss of these traits from the gene pool is irreversible and limits the capacity of this species to resist a changing climate – and that of breeders to produce more resilient varieties. An understanding of the evolution of *O. glaberrima* and its adaptation to different natural environments is therefore an important step in characterising the agronomic potential of this species, the protection of which will be indispensable for sustaining genetic crop diversity and a food secure future.

### Evolutionary origins

Two main competing hypotheses have been proposed concerning the domestication of rice in Africa. One proposes that plant domestication in Africa occurred in a non-centric manner, over a protracted period of time [8], and has been called the ‘protracted transition model’. According to this hypothesis, rice was domesticated in multiple areas of domestication in West Africa, without a defined moment and centre of origin [8]. The other proposes a single centre of domestication along the Niger River, followed by two secondary diversification events: one along the coast of what are now the countries of Senegal and Gambia, and one in the Guinea Highlands [2]. This has been called the ‘rapid transition model’. According to a particular theory supporting the latter hypothesis, domestication was triggered at an acute time point when climate change started transforming forests into savannah around 4000 years ago [9]. The sudden drought meant that the increasing population could no longer rely on traditional forest products. However, the nature of hunter-gatherer and pastoral societies in West Africa calls into question whether a definite centre of origin is likely ever to be found [10]. Human migration may have assisted the exchange of particular rice varieties between ethnic groups, diminishing the differences between them. In addition, ongoing hybridisation with *O. barthii* and later cultivation alongside *O. sativa* may have caused interspecific gene flow, which further complicates inferences about domestication origin [11].

In addition to the centric and non-centric model, an additional theory about crop domestication stipulates multiple (usually two) defined centres, as has been observed in both Asian rice [12] and barley [13]. Such a polycentric origin can explain the existence of two distinct, geographically separated sub-populations or even sub-species, like *O. sativa* ssp. *indica* (which originated in India) and *O. sativa* ssp. *japonica* (which originated in China). These subspecies of rice have separate origins, although later domestication stages saw extensive gene flow between the two, which has been associated with the transfer of domestication alleles [14]. An overview of the various domestication hypotheses is presented in Fig 1.

**Fig 1.**
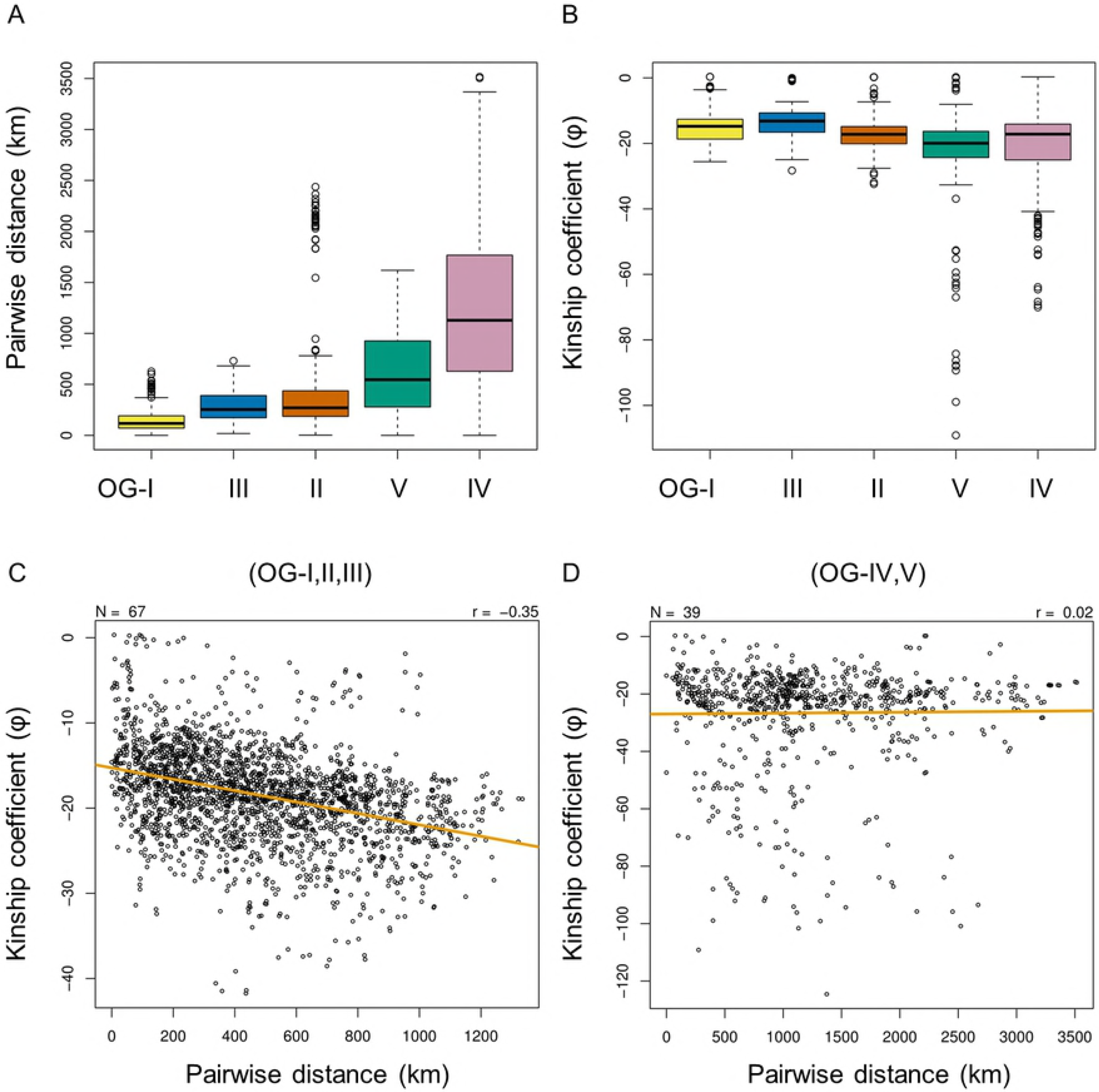
Overview of crop domestication hypotheses. The debate about African plant domestication has mainly revolved around the non-centric model and centric model. An alternative model is derived from the idea of domestication centres but holds that there can be more than one.

### Genetic characteristics

Several studies have tried to illuminate the question of how and where African rice originated, either implicitly or explicitly addressing the domestication hypotheses described above. A study of 14 unlinked nuclear genes in 40 individuals found *O. glaberrima* to have 70% lower genetic diversity and hardly any population structure compared to its wild relative, supporting a single origin around the Inner Niger Delta [15]. In contrast, Semon et al. [16] found five cryptic sub-populations based on a study of 93 microsatellite markers in 198 *O. glaberrima* and 9 *O. sativa* accessions. Two of the sub-populations showing interspecific admixture with *O. sativa* were later shown to be true *O. sativa* varieties [11]. The three other groups were specific to *O. glaberrima* and associated with different phenotypic traits, corresponding to the floating, non-floating and upland ecotypes, respectively [16]. The authors found no isolation by distance and therefore argued that the maintenance of sub-populations was mainly caused by artificial selection and human-mediated gene flow. A study of genetic diversity based on 235 single nucleotide polymorphisms (SNPs) also found two distinct populations in 266 *O. glaberrima* accessions without any correlation to geography – in contrast to their wild relative, which clustered into three geographically distinct sub-populations [11]. Contrary to earlier findings, however, this study failed to link population structure to phenotypic traits.

A potential shortcoming of these earlier studies is the type and quantity of genetic markers analysed, which leads to a low resolution of genetic diversity. This limitation is overcome with the introduction of genome-wide analyses and tools. Nabholz et al. [17] analysed more than 12,000 RNA transcripts from 10 wild and 9 domesticated African rice individuals and thus confirmed that *O. glaberrima* underwent an extreme genetic bottleneck and is probably the least genetically diverse crop ever documented. This low level of diversity was confirmed by subsequent genomic studies. In 2014, Wang et al. [18] published an assembly of the currently only available reference genome of *O. glaberrima* (AGI1.1). Whole genome resequencing of 94 *O. barthii* and 20 *O. glaberrima* accessions revealed that domesticated rice consistently clustered with one of five sub-populations of wild rice, suggesting a single origin in the area of Senegal, Gambia, Guinea and Sierra Leone [18]. A genome-wide association study of 93 additional *O. glaberrima* accessions fine-tuned this finding by providing evidence for geographically localised diversification within this region, specifically suggesting that reduced salt tolerance may have evolved in the tropical south of the western Atlantic coast in response to higher rainfall and reduced salinity [19]. In addition, this study supports a period of low-intensity cultivation that may have started as early as 10,000 years before the effective population size reached a low point around 3000 years ago, when African rice was reputedly domesticated.

### One or multiple origins?

Thus far, no conclusive evidence has been provided in favour of either the non-centric or the centric domestication hypothesis. Although the first genome study of African rice supported a single origin of domestication [18], the suggested place of origin is not located along the Inner Niger Delta as suggested by Portères [2], but rather in what Portères proposes to be the secondary diversification centre(s) on the Atlantic coast. In contrast, the competing scenario of diversification in this region makes no claims as to the original centre of domestication, and even provide evidence for the protracted model [19]. None of the other genetic studies mentioned in the previous section were able to pinpoint a clear centre of domestication. In addition, evidence regarding population structure is inconclusive and varies from no observed structure at all [15,17], to clearly differentiated [11,16] and geographically localised [19] sub-populations. Furthermore, the use of widely divergent types and quantities of data, including RNA transcripts, microsatellites, gene markers and genome-wide SNPs, precludes a systematic comparison of the results of these studies.

The available genomic data enable a reinterpretation of previous results and might clarify some of the present ambiguities regarding the origin and diversification of African rice. Thus, while many questions concerning the domestication and migration of African rice are still outstanding, the complete genome sequences of more than a hundred *O. glaberrima* accessions – and an almost equal number of *O. barthii* accessions – provide a wealth of genetic data that can be used to reconstruct the evolutionary history of African rice, and to compare the genetic diversity of *O. glaberrima* in its different localities.

In order to elucidate the origin and diversification of *O. glaberrima* in West Africa, we performed a critical reassessment of the publicly available whole genome resequencing data of *O. glaberrima* and *O. barthii* accessions from across the species range (S1 Table and S2 Fig). These data were mapped against the *O. glaberrima* reference genome and analysed through a combination of population genetic and phylogenetic approaches, details of which can be found in the Materials and methods. In short, we wanted to 1) confirm the genetic bottleneck in African rice as a result of domestication; 2) identify which alleles have been driven to (near) fixation as a result of artificial selection; 3) discover population structure and differentiation within and between the two species; 4) assess the influence of geography on the distribution of genetic variation; 5) explore discordances between the evolutionary histories of candidate domestication genes and the genome-wide species tree; and 6) predict the functional relevance of gene regions with divergent histories.

The first objective was met by quantifying the relative genetic diversity of *O. glaberrima* and *O. barthii* through a variety of statistics. The second objective was met by unfolding the site frequency spectrum and performing a selection scan. The third objective was met by estimating the number of ancestral populations to delimit extant populations and calculating the fixation index between them. The fourth objective was met by mapping a principal components analysis onto known geographic coordinates and measuring isolation by distance. The fifth objective was met by phasing gene haplotypes and comparing their relationships with the overall phylogeny based on genomic distances. Finally, the sixth objective was met by identifying high impact mutations based on computational analyses and previously published experimental results. These results were subsequently used to weigh the evidence for local versus global adaptation, discuss the taxonomic implications for species delimitation, argue for a single or multiple domestication events, and ultimately to reconsider the dominant domestication hypotheses.

## Results

### Genetic bottleneck

Joint variant calling and quality filtering resulted in a total of 3,923,601 SNPs. Average SNP density is almost twice as high in *O. barthii* (9.04 SNPS per kb) as in *O. glaberrima* (5.00 SNPs per kb). About half of the polymorphic sites in *O. barthii* are unique to this species. In contrast, the vast majority of polymorphic sites in *O. glaberrima* is shared with *O. barthii*, suggesting very little species-specific variation among the domesticated accessions. The ratio of synonymous to non-synonymous substitutions and the relative portion of protein coding variation appear to be roughly the same (between 0.80-0.85, and less than 10% of the total SNP count, respectively). Both *O. barthii* and *O. glaberrima* are predominantly selfing plants, which is reflected in their low levels of heterozygosity. Removal of low coverage individuals (< 4X) in both species revealed an even stronger reduction of heterozygosity in *O. glaberrima*, consistent with a higher level of inbreeding.

Relative nucleotide diversity between the two species was significantly lower in the cultivated species (π_c_ = 0.0007) than in the wild species (π_w_ = 0.0013) at p < 1.0E-05. The relative nucleotide diversity (π_w_/π_c_) was found to be 1.87 across the genome, but was markedly higher in some regions (S3 Fig). Tajima’s D was significantly different between the two species at p < 1.0E-05, being predominantly negative in *O. glaberrima* (−0.6761) and positive in *O. barthii* (0.5172). The relative levels of Tajima’s D suggest that large parts of the *O. glaberrima* genome exhibit an excess of rare variants as compared to *O. barthii*. This is compatible with the general trend in π ratio (π_w_/ π_c_), which is usually well above one (S3 Fig) and takes on particularly high values when Tajima’s D is extremely negative.

The excess of rare variant in *O. glaberrima* is confirmed by the Minor Allele Frequency (MAF) spectrum (S3 Fig), where *O. glaberrima* has a larger spike in low frequency alleles (MAF < 0.01) than the majority of *O. barthii* alleles, which are of intermediate frequency (MAF 0.01-0.05). The low levels of nucleotide diversity and the large number of rare variants found in *O. glaberrima* are consistent with a scenario of population expansion following a sudden drop in effective population size. These results are in congruence with previous findings [17–19] and indicate that a strong reduction in diversity in *O. glaberrima* occurred as a result of domestication.

### Artificial selection

The previous statistics show a deviation from neutrality that could be caused both by changes in the effective population size as well as selection. Whereas demographic factors can usually account for extremely low frequencies of derived alleles, it is very unlikely that genetic drift alone can push derived alleles to extremely high frequencies. A U-shaped derived allele frequency spectrum is therefore used as evidence of positive selection, but has not been demonstrated in African rice to date.

In contrast to the expected site frequency spectrum under neutral conditions, a large number of high frequency derived alleles is observed in both species (Fig 2). Despite this excess of high frequency derived alleles, *O. glaberrima* shows a greater excess (35% of total SNPs above expectation) than *O. barthii* (27% of total SNPs above expectation) in the far-end of the spectrum (0.7 – 1.0). This excess is also caused by higher frequency classes in *O. glaberrima* (21% of total SNPs > 0.99) than in *O. barthii* (18% of total SNPs > 0.95). Over the whole spectrum, the discrepancy between observed and expected frequencies as observed in an empirical cumulative distribution plot, is greater in *O. glaberrima* than in *O. barthii* (Fig 2). This difference was found to be significant in a two-sample Kolmogorov-Smirnov test (p < 0.5E-04). The disproportional skew in favour of high frequency derived alleles in *O. glaberrima* suggests that large parts of the genome bear signs of recent positive selection.

**Fig 2.**
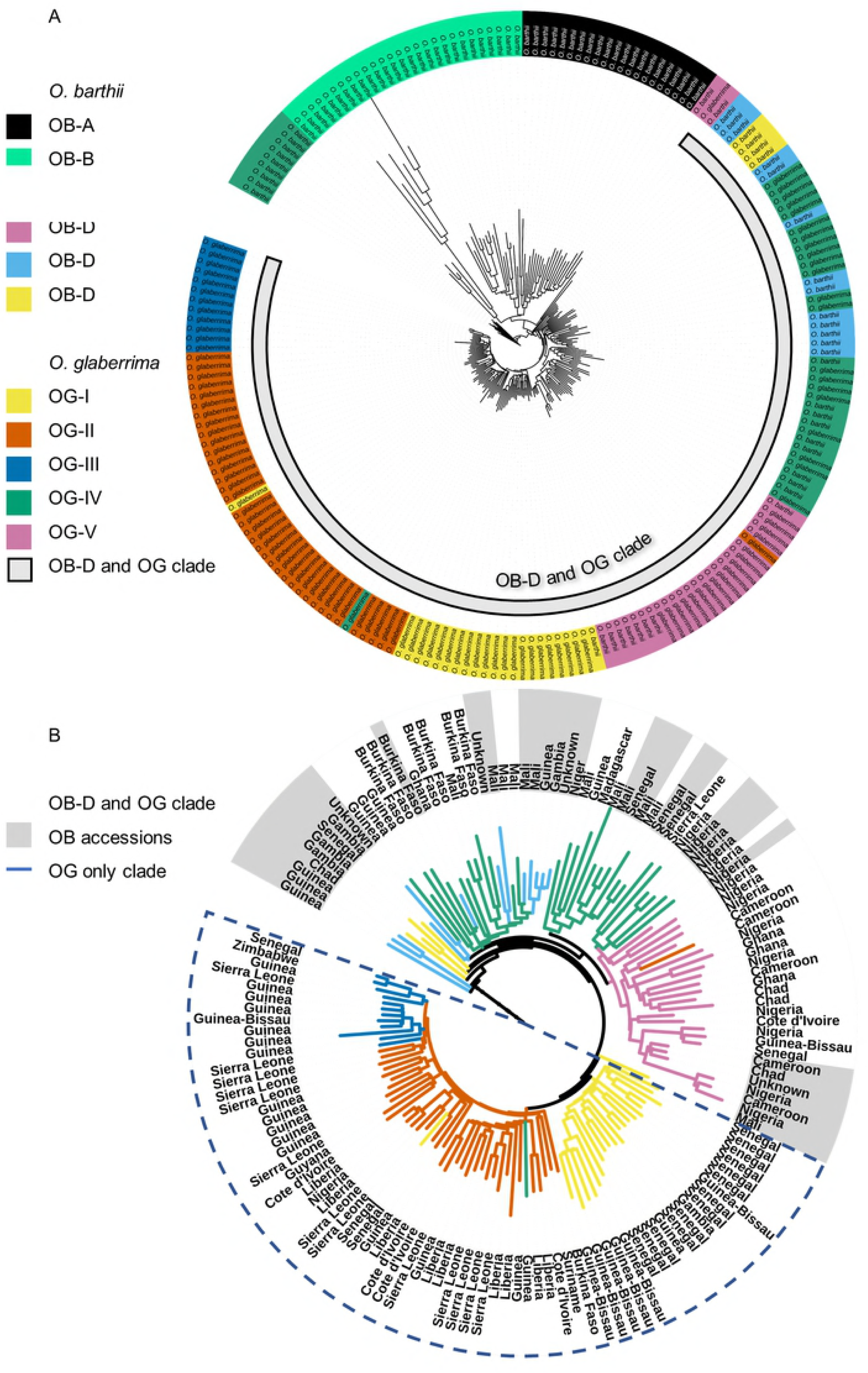
Derived allele frequency spectrum of non-coding and synonymous substitutions in *O. barthii* and *O. glaberrima*. A. Observed and expected marginal derived allele frequency spectrum of *O. glaberrima*. B. Observed and expected marginal derived allele frequency spectra of *O. barthii*. C. Derived allele frequency spectra of *O. glaberrima* and *O. barthii* together. D. Empirical cumulative distribution function of the difference between the expected and observed frequency spectra.

To further investigate the regions of the genome that might have been under recent and strong selection, a composite likelihood ratio (CLR) test was conducted (S4 Fig). Overall, the genome-wide CLR was higher in *O. glaberrima* (1.12 on average) than in *O. barthii* (0.93 on average). Barring a single shared outlier on chromosome 4, there is a remarkable lack of overlap of outliers between the two species, suggesting that the sweeps found in *O. glaberrima* are unique to the domesticated accessions. More surprising, however, is the fact that out of 20 candidate domestication genes, not a single one shows clear-cut evidence of a recent, strong selective sweep (S5 Fig). This is either because the chosen selection scan has problems separating the effects of demography from those of selection, or because the model of a single, hard sweep fails to explain the history of these genes. If the latter is the case, this calls into question the ‘single origin’ hypothesis and the domestication of *O. glaberrima* might have resulted from more complex processes than simple selection scans are able to detect. Since these scans presume that a variant under selection swept through an entire population, the possibility that part of the population escaped the sweep, either due to different selection pressures or due to population substructure, remains unexplored.

### Ancestral and extant populations

To investigate to what extent population substructure in *O. glaberrima* and *O. barthii* could disqualify the hard sweep model, we re-examined the variation in both species. A joint ADMIXTURE analysis was performed to infer which ancestor fractions are shared between the species and which are unique. Cross-validation error estimates shows that model fit was optimised for both species at K=8 and at K=5 when either *O. glaberrima* or *O. barthii* was considered by themselves (S6 Fig). Two of the eight ancestral populations were predominantly found in *O. barthii* alone, and three others almost exclusively in *O. glaberrima* (Fig 3A).

**Fig 3.**
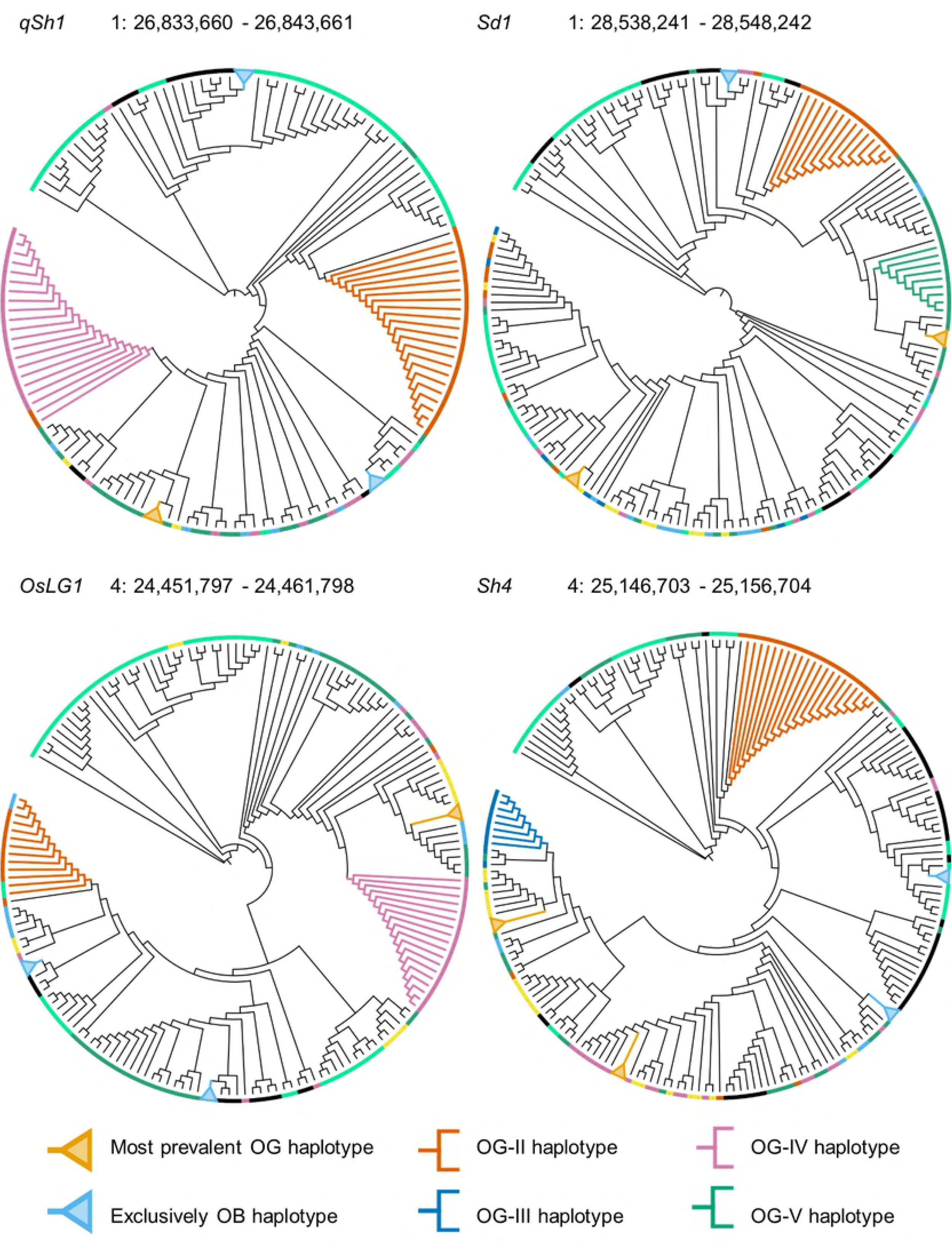
Structure and geographic clustering of genetic variation in *O. barthii* and *O. glaberrima*. A. Admixture analysis with K = 8 ancestral populations of wild and domesticated rice accessions combined. B. Admixture analysis with K = 5 ancestral populations of domesticated rice accessions only and their corresponding collections sites. Accessions were assigned to a cluster according to their genetic background, with the colour of each sample representing the ancestral population (K = 5) that accounted for the majority (>50%) of pruned SNPs in that accession. C. PCA of all geo-referenced accessions of *O. glaberrima* (dots, filled) and *O. barthii* (triangles, open) together. The first principal component is correlated significantly with latitude and the second principal component is correlated significantly with longitude. D. PCA of all geo-referenced *O. glaberrima* accessions separately. The first principal component is correlated significantly with longitude and the second principal component is correlated significantly with latitude.

Based on these results, the two species were subdivided in roughly evenly sized sub-populations, depending on the dominant ancestor fraction in each individual: *O. barthii* in sub-populations OB-A through OB-D, and *O. glaberrima* in OG-I through OG-V, respectively (Fig 3). Interestingly, OB-C and OB-D represent the individuals that were previously found to form a clade with *O. glaberrima* [18]. This makes sense in light of the observation that these individuals contain ancestor fractions that are also found in *O. glaberrima*, in contrast to the individuals from populations OB-A and OB-B. The only domesticated populations that do not appear to share ancestry with any of the wild populations, are OG-II and OG-III. A large number of individuals in these populations contain substantial fractions of both ancestries, making these populations less readily distinguishable.

The collection sites of OG accessions suggest that the observed population structure has a strong geographic component (Fig 3B). Whereas most of the coastal accessions belong to either OG-I, OG-II or OG-III, the majority if accessions sampled inland belong to OG-IV and OG-V. Geographic populations of *O. glaberrima* have been proposed before by Meyer et al. [19], who separated African rice into groups based on a 11°N and the 6°W cline. Association of the coordinates of collection sites with genetic variation was tested on the basis of a principal component analysis (PCA), conducted for both species together and separately. We were unable to detect any such association in *O. barthii* alone. This may be due to the low number of geo-referenced individuals, a lack of geographic structure or both. When both species were pooled together (Fig 3C) or *O. glaberrima* was taken alone (Fig 3D), however, latitude and longitude were found to be significantly correlated with either of the top two principal components. This confirms that genetic variation is partially explained by the site of collection, at least in the domesticated species.

### Geographic distribution of genetic variation

To determine the degree of genetic differentiation between the geographic regions, the fixation index (F_ST_) was calculated between individuals separated by the 11°N and the 6°W cline, respectively. Genetic differentiation between *O. glaberrima* and *O. barthii* was calculated as a reference point for within *O. glaberrima* comparisons (Table 1). While *O. glaberrima* and *O. barthii* are differentiated greatly (F_ST_ > 0.15), the western and eastern groups were differentiated moderately (F_ST_ = 0.10) and the northern and southern groups were differentiated only a by a small degree (F_ST_ < 0.05). This is in line with previous results, proposing an early split between the eastern and western cultivation range, followed by a later split between the northern and southern diversification centres [19].

**Table 1.**
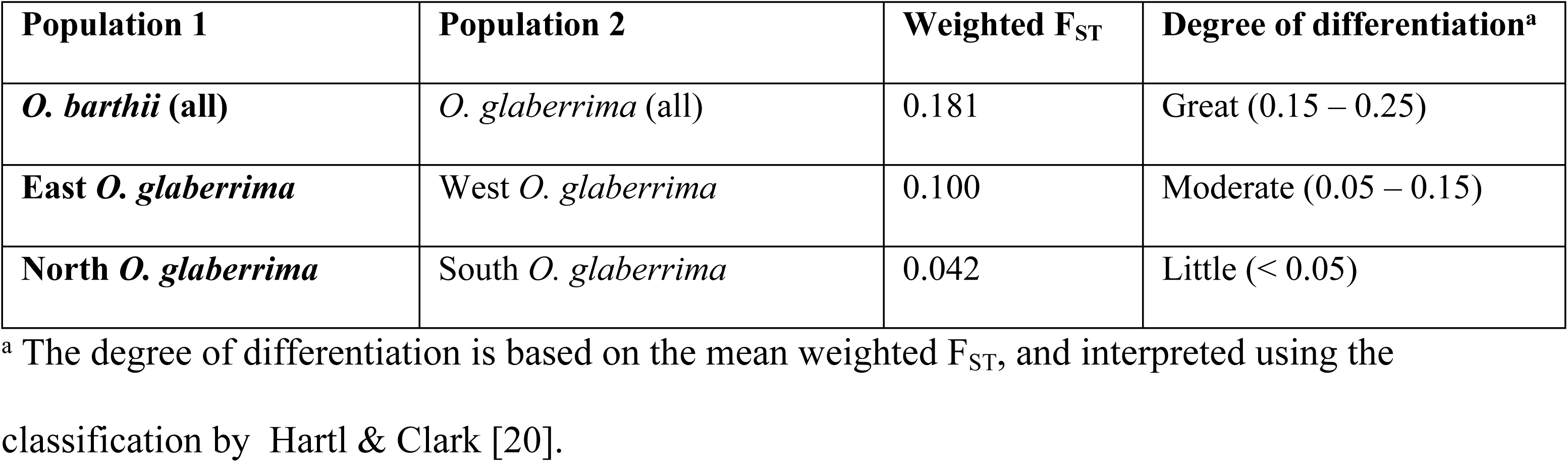
Fixation index (F_ST_) between species and between geographic clusters.

Genetic differentiation from *O. barthii* for all five *O. glaberrima* populations was found to be large (F_ST_ > 0.15), but relatively larger for the three coastal populations (F_ST_ > 0.25) than for the two inland populations (F_ST_ 0.15 – 0.25). Genetic differentiation from *O. barthii* is the smallest for OG-IV and the largest for OG-II (Table 2). This pattern is mirrored by the number of segregating sites remaining in each population after removing monomorphic SNPs, which is again the largest for OG-II and the smallest for OG-IV (Table 2). The opposite trend can be seen for average nucleotide diversity, which is smaller in the coastal populations (π < 1.0/kb) than in the inland populations (π > 1.0/kb). We thus observe that, even though the total number of polymorphic sites in larger in OG-I through OG-III, the average number of pairwise differences between these individuals is lower. This suggests that a smaller number of individuals carries a larger fraction of the polymorphic sites, which is consistent with a population expansion scenario, as explained before.

**Table 2.** Genetic attributes of five genetic *O. glaberrima* populations. Columns show relative sample size, degree of polymorphism and genetic differentiation from *O. barthii*. Fixation index (F_ST_) reflects the differentiation between a subset of individuals (n = 15) from each population and an equal number of *O. barthii*.

| Population | Accessions | Segregating sites | $\pi/\text{kb}$ | Weighted $F_{ST}$ | Degree of differentiation<sup>a</sup> |
| --- | --- | --- | --- | --- | --- |
| OG-I | 22 | 1,443,290 | 0.9905 | 0.25641 | Very great ($> 0.25$) |
| OG-II | 30 | 1,550,660 | 0.7504 | 0.30894 | Very great ($> 0.25$) |
| OG-III | 19 | 1,646,372 | 0.9166 | 0.27588 | Very great ($> 0.25$) |
| OG-IV | 21 | 1,236,010 | 1.179 | 0.20514 | Great (0.15 – 0.25) |
| OG-V | 20 | 1,324,300 | 1.076 | 0.24287 | Great (0.15 – 0.25) |

<sup>a</sup> The degree of differentiation is based on the mean weighted $F_{ST}$ , and interpreted using the classification by Hartl & Clark [20].

The combined evidence of the previous sections suggests that the increase in genetic differentiation from the inland to the coastal populations may be linked to geographic range expansion. To test whether the observed population structure could be the result of geography, isolation by distance (IBD) was assessed among all West African accessions. Genetic IBD is explained by the accumulation of genetic differences by dispersal [21]. We evaluated genetic IBD as the correlation between pairwise relatedness, measured by the kinship coefficient (φ), and pairwise geographic distance, measured by the shortest distance between the collection sites of two accessions in kilometres. Genetic IBD was discernible among the coastal populations with a correlation coefficient (*r*) of −0.35 (Fig 4). This correlation was stronger than the observed correlation within any single population, or in all populations pooled together (S7 Fig). In contrast, there was hardly any IBD among the inland populations, with a correlation coefficient (*r*) of only 0.02 (Fig 4F). This suggests that, whereas in the inland regions geographic distance seems to be a very poor indicator of relatedness, some of the population structure observed along the coast can indeed be explained by geographic distance. This would correspond to the accrual of mutations as *O. glaberrima* dispersed throughout the coastal range.

**Fig 4.**
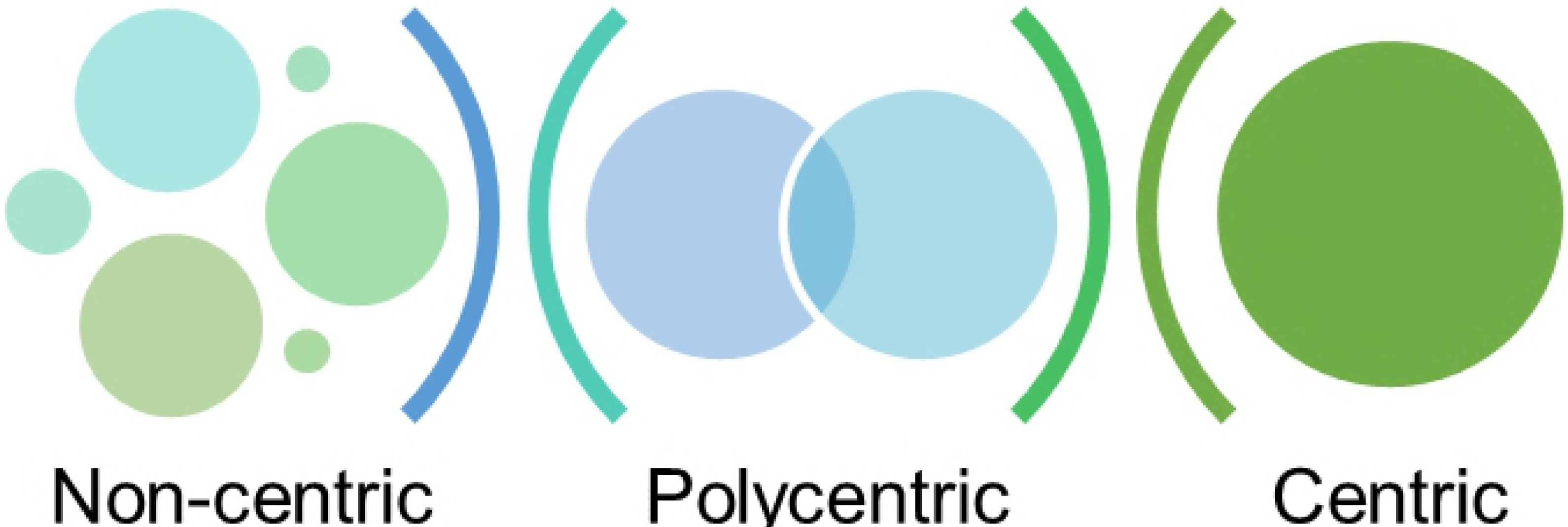
Genetic isolation by distance in *O. glaberrima.* A. Distribution of the pairwise geographic distances between individuals in kilometres, grouped per population. B. Distribution of the kinship coefficients between individuals, grouped per population. C. Isolation by distance among the coastal populations (OG-I, OG-II and OG-III). Outliers (separated by more than 1500 km) were omitted. D. Isolation by distance among the inland populations (OG-IV and OG-V). Outliers (separated by more than 3500 km) were omitted. Each dot symbolises a unique pair of individuals within in each group. Whereas N denotes the number of accessions included in each analysis, the number of pairwise comparison equals N! and is therefore markedly higher.

### Whole-genome and candidate gene trees

The progressive differentiation of *O. glaberrima* from *O. barthii* in an east to west direction, is reflected in a genome-wide neighbour-joining (NJ) tree of all accessions (Fig 5). Almost all the coastal accessions (OG-I, OG-II and OG-III) appear to form a clade that do not include any *O. barthii* or any of the inland accessions (OG-IV an OG-V). In contrast, the closest wild relatives of *O. glaberrima* (OB-C and OB-B) cluster primarily among the inland accessions, whereas OB-A and OB-B together form a monophyletic clade with much longer branches than the other *O. barthii*. This clustering of *O. glaberrima* and *O. barthii* populations is largely confirmed by PCA and multi-dimensional scaling of the genetic variation in both species, where the distinct nature of OB-A and OB-B is visible in its large distance from the other accessions.

**Fig 5.**
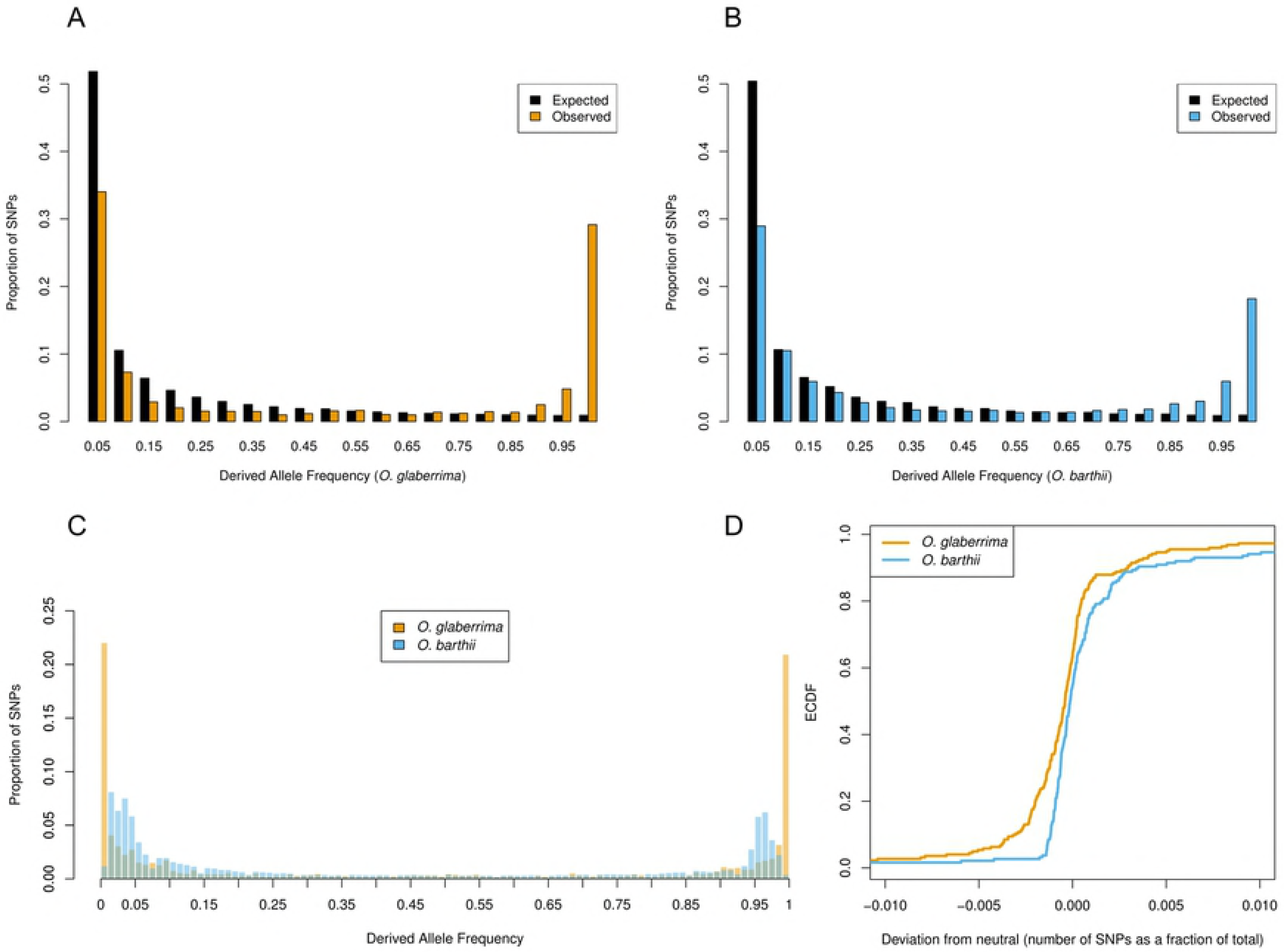
Neighbour Joining (NJ) tree of *O. barthii* and *O. glaberrima* whole genome sequences, based on 3,923,601 genome-wide SNPs. A. NJ tree with all *O. barthii* (OB) and *O. glaberrima* (OG) accessions. Accessions are labelled by species and coloured according to their genetic cluster (K = 8). The grey bar labelled ‘OB-V and OG’ indicates the smallest monophyletic clade containing all *O. glaberrima* and its nearest wild relatives. B. Pruned NJ tree with only OB-V and OG accessions, representing the smallest monophyletic clade that contains all *O. glaberrima* and its nearest wild relatives. Branches are coloured according to genetic cluster (K = 8) and are labelled by country. OB-V accessions labels have a grey background. The dashed blue line surrounds the largest clade that contains only OG and no wild relatives

The clustering of inland samples with *O. barthii* implies that they share a common pool of genetic variation from which *O. glaberrima* was domesticated, and that the coastal lineages branched off at a later point in time. The smaller nucleotide diversity in the coastal populations, indicative of recovery from a genetic bottleneck, seems to support this scenario. Assuming that the sampling locations of the present accessions reflect their historical origins, this suggests that the origin of domestication lies east of the 6°W cline, and that *O. glaberrima* subsequently migrated westward. This is consistent with the TreeMix analysis performed by Meyer et al. [19], and with the domestication hypothesis proposed by Portères [2].

To assess whether certain domestication traits of *O. glaberrima* could have had multiple origins, NJ trees were constructed for several domestication genes known from recent rice genetics literature [18,22], a list of which can be found in S8 Table. We labelled the five most common haplotypes per gene and then annotated the trees based on population structure, to see which of the *O. glaberrima* subpopulations segregate into different haplotypes and whether they cluster with the expected OB-C and OB-D accessions.

Eight of the twenty genes clearly deviate from the genome-wide phylogenetic signal (S9 Table). In these genes, a subset of *O. glaberrima* from a single subpopulation cluster together in smaller clades that are far removed from the larger *O. glaberrima* clade. In four of those (*Sd1, qSh1, OsLG1* and *Sh4*), the segregating haplotype that is farthest removed from the major clade in these genes is composed *O. glaberrima* individuals that are exclusively from the OG-II population (Table 3). A closer inspection of the neighbouring *O. barthii* accessions reveals that their closest relatives all belong to the OB-B subpopulation, rather than the expected OB-C and OB-D populations (Fig 6). A re-examination of the other trees subsequently shows that some of the OG-II accessions also cluster with OB-B in other genes (Table 3), although their numbers did not warrant their inclusion as one of the five largest haplotypes. This recurrent pattern stands in stark contrast with the genome-wide phylogeny, where the OB-B population is genetically most distant from *O. glaberrima* (Fig 5).

**Table 3.** OG-II accessions possessing a different haplotype than the main *O. glaberrima* clade in multiple domestication genes. All these accessions have in common that they cluster with OB-B rather than OB-C and OB-D and share a single haplotype with other *O. glaberrima* individuals for the genes that are marked with an ‘x’.

| Accession | *qSh1* | *Sd1* | *OsNAC6* | *OsLG1* | *Sh4* | *GW2* | *COLD1* | *Phr1* | *Rc* | Country |
| --- | --- | --- | --- | --- | --- | --- | --- | --- | --- | --- |
| IRGC103937 | x | x | x | x | x |  |  |  |  | Liberia |
| IRGC103946 | x |  |  |  | x |  |  |  |  | Liberia |
| IRGC103949 | x | x | x |  | x |  |  |  |  | Liberia |
| IRGC103953 | x | x | x | x | x |  |  |  |  | Sierra Leone |
| IRGC103988 | x | x | x |  | x |  |  |  |  | Sierra Leone |
| IRGC104035 | x | x | x | x | x |  |  |  |  | Cote d'Ivoire |
| IRGC104036 | x | x | x | x | x |  |  |  |  | Cote d'Ivoire |
| IRGC104165 | x | x | x |  | x |  |  |  |  | Guinea |
| IRGC105048 | x |  |  |  | x |  | x | x | x | Liberia |
| IRGC105049 | x |  |  |  | x | x | x |  |  | Liberia |
| TOG6203 | x | x | x | x | x |  |  |  |  | Guinea |

**Fig 6.**
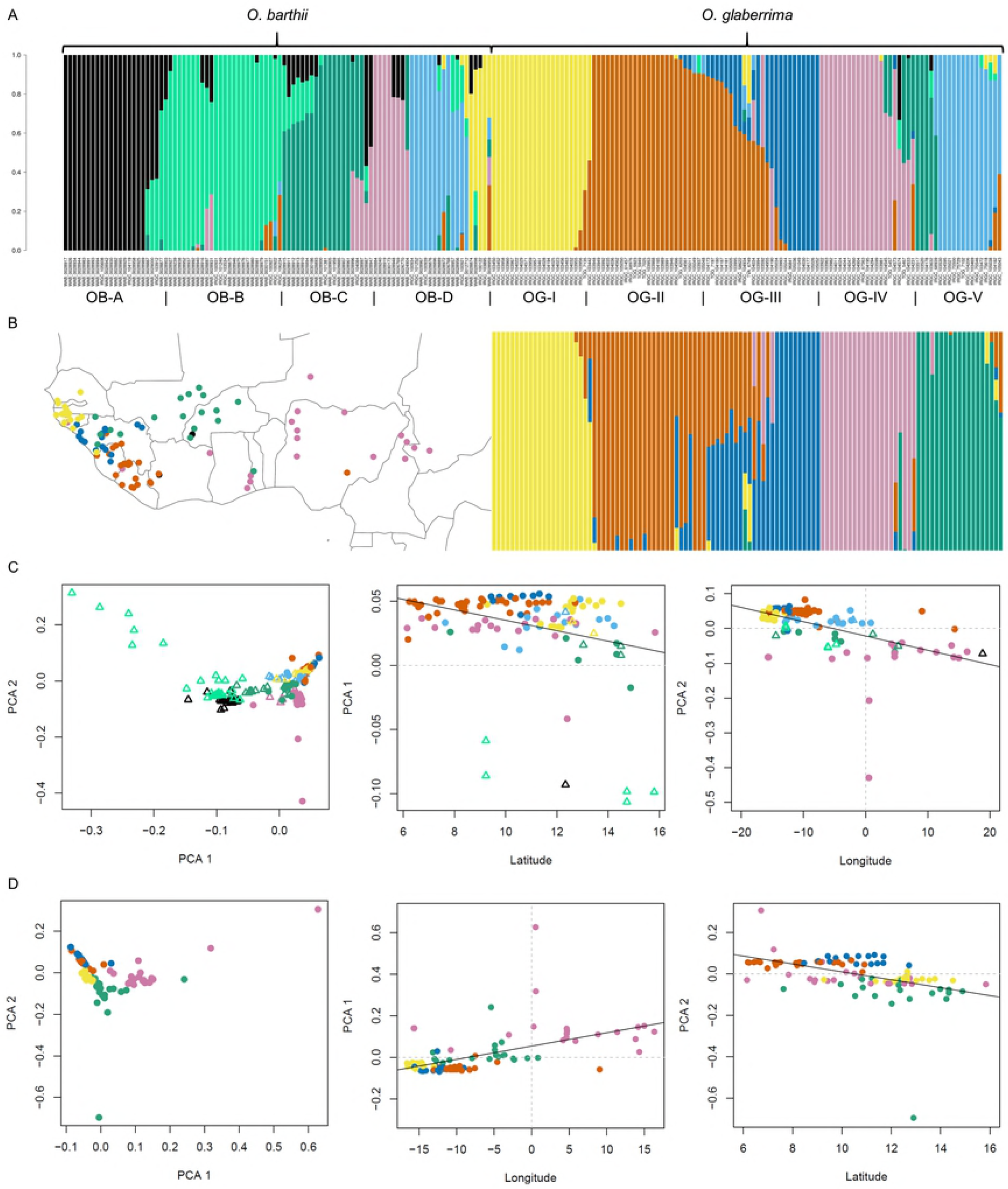
Separation of OG-II haplotype in multiple domestication genes. The five (or in case of equal haplotype counts, six) largest haplotypes are labelled. The most common haplotypes, containing accessions from multiple sub-populations *O. glaberrima*, are collapsed into orange nodes. Haplotypes that consist exclusively of *O. barthii* are collapsed into blue nodes. Remaining haplotypes, consisting of a mix *O. barthii* and *O. glaberrima* accessions from a single subpopulation, are expanded with branch colours reflecting their population of origin of the *O. glaberrima* accessions

Haplotypes restricted to a single subpopulation are also observable in a number of other genes (S10 Fig). Most notably, accessions from the OG-IV subpopulation do not only cluster separately in *qSh1* and *OsLG1*, but are also part of smaller haplotypes in *Phr1*, *MOC1*, *Rc* and *Ipa1* (Table 4). Although these accessions are exclusively surrounded by *O. barthii* accessions sharing the same ancestry, the fact that they frequently form a distant clade provides further support for the separate roots of domestication in the coastal and inland regions of West Africa.

**Table 4.**
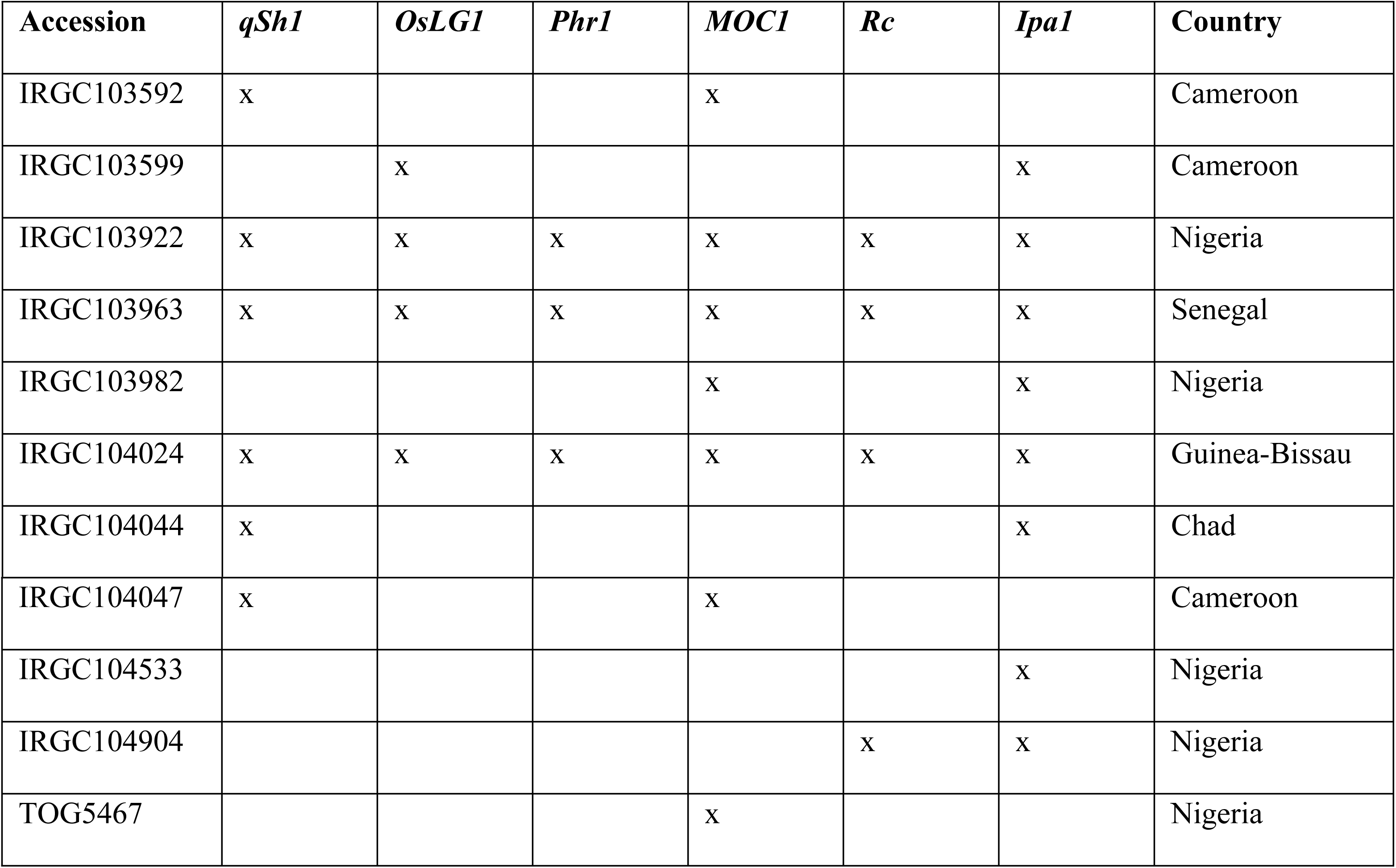
OG-IV accessions possessing a different haplotype than the main *O. glaberrima* clade in multiple domestication genes. All these accessions have in common that they cluster with *O. barthii* accessions of similar ancestry and share a single haplotype with other *O. glaberrima* individuals for the genes that are marked with an ‘x’

### Functional characteristics of haplotypes

To see what phenotypes might be associated with the segregating haplotypes in these genes, we evaluated the impact of the responsible substitutions using variant prediction software. No high-impact substitutions were detected apart from a variant in *Sh4* (a shattering gene), which encodes a premature stop codon in almost all domesticated accessions. Polarisation of this SNP against *O. meridionalis* as an outgroup demonstrated that the variant responsible for truncation of the protein is derived, and the variant encoding the intact protein is ancestral. The ancestral variant was found to be represented exclusively in the OG-II haplotype. Since the non-shattering phenotype is a crucial trait in the domestication syndrome, the ancestral state of this substitution in a limited number of OG-II accessions presupposes that another variant – either in the same gene or in a different gene – might be causing the same phenotype. This has recently been confirmed by functional characterisation of another gene, called *Sh3*, that is on its own responsible for a non-shattering phenotype in African rice [23].

Although the functional relevance of these genetic differences has been demonstrated for one gene (*Sh4*), both *in silico* and *in vitro*, the phenotypic consequences of differentiation in the other genes remain to be determined. Considering the extensive LD in *O. glaberrima* and the fact that strong candidates for high impact substitutions could not be identified, it cannot be excluded that associated functional mutations lie outside the intervals included in our phylogenetic analyses and that the gene haplotypes identified here may have hitchhiked on the selection of a different genomic feature altogether. Any claims regarding the adaptive significance of these phylogenetic patterns will therefore have to be complemented by experimental evidence.

Interestingly, screening of homologs of additional agronomically important genes reveals that the NAC transcription factor *OsNAC6* is located less than 15kb away from *Sd1*. *OsNAC6* has been identified as a key regulator of rice stress responses and has been shown to enhance drought and salinity tolerance [24]. Phylogenetic analysis shows that the exact same OG-II accessions form a separate haplotype in this gene as in *Sd1* (Table 3). A genome-wide association study of *O. glaberrima* has provided evidence for geographical divergence of salt tolerance traits in the SW coastal population and suggested two other candidate genes *OsHAK5* and *OsHAK6* that were linked to a significant SNP on the far-end (position 30698514) of chromosome 1 [19]. This gives further credibility to the idea that the segregating haplotypes in these genomic regions may underlie functional differentiation of the coastal OG-II accessions.

## Discussion

### Modes of adaptation

This is the first time that the data sets of two large scale genomic studies on African rice [18,19] have been combined and comprehensively reanalysed, allowing for several important new insights into the domestication history of *O. glaberrima*. The sample sizes of both *O. glaberrima* and *O. barthii*, in combination with the genomic nature of the data, confer an advantage to the analyses presented here that previous studies did not have.

The diversity measures reported in this study are consistent with previous reports [18,19] and provide strong evidence for a large reduction in diversity in the genome of *O. glaberrima* as a result of domestication. This reduction in diversity is most likely caused by a combination of selection, favouring a small number of preferred alleles, and demographic history, causing a large drop in effective population size (*N*^*e*^).

Both the negative values of Tajima’s D and the skewed MAF distribution in favour of rare alleles support the notion that *O. glaberrima* underwent population expansion following a severe bottleneck. While this demographic scenario can account for the large number of rare derived alleles, it is unlikely to account for the large number rare ancestral alleles. The fact that *O. glaberrima* shows a larger excess of high frequency derived alleles than *O. barthii*, as evidenced by their empirical cumulative distribution functions, is an indication that it at least underwent stronger positive selection than its wild relative.

Despite this evidence, the identification of exact regions in the genome that have been under positive selection is notoriously difficult due to the confounding effects of demographic history, which are known to produce local reductions in genetic diversity that can look remarkably like selective sweeps [25]. None of the currently available selection scans have been tested for their robustness under simulations of a very extreme and rapid bottleneck, as encountered in domestication events. Without additional modelling, therefore, we cannot be sure that demographic history of African rice did not interfere with the CLR test presented here. In addition, this test does not consider alternative forms of selection, such as balancing selection or diversifying selection. In recent years, more studies have focused on the existence of ‘soft’ selective sweeps, which are caused by positive selection on either pre-existing variants or multiple de novo mutation. we did not explore alternative models of selection in this study, because the existence of several haplotypes due to multiple novel mutations or standing variation renders the detection of soft sweeps exceedingly complex.

Perhaps the most compelling argument that can be made against the hard sweep model in *O. glaberrima*, is the population subdivision. A recent study of the AfricaRice gene bank collection also revealed exactly five genetic clusters based on a study of 27,560 SNPs across 2,179 accessions. These clusters are linked strongly to country of origin, but less so to ecotype [26].

The geographic component of population structure can prevent instances of position selection from sweeping through the entire population in multiple ways. Firstly, isolation by distance delays the migration of a beneficial allele, thus diminishing the effect of genetic hitchhiking that is observed in a hard sweep [27]. In addition, population structure can cause parallel adaptation to a global selection pressure in geographically separated demes [28], resulting in a soft sweep rather than a hard sweep. Geographically separated populations may also undergo local adaptation due to geographically localised selection. This has been observed in the case of drought tolerance in the coastal populations of *O. glaberrima* [19]. Hence, it is not unlikely that population substructure further complicates the detection of sites that are universally under positive selection throughout the entire species.

### Species delimitation

The population structure of African rice further sheds light on the geography of its domestication. Interestingly, all the accessions that originate from the proposed primary domestication centre of African rice, around the Inner Niger Delta, belong to a single genetic cluster (OG-V). This cluster is also found in Guinea, where it coincides with one of the coastal populations (OG-III) and where the species splits into two other populations: OG-I to the north and OG-II to the south. The geographic ranges of the latter populations roughly correspond with the proposed secondary domestication centres in what used to be Senegambia and in the Guinea Highlands, respectively. In contrast, OG-IV is restricted to the inland areas and is located primarily south and east of OG-V. The fact that OG-IV and OG-V appear to be the most genetically diverse and least genetically differentiated from *O. barthii*, while the reverse can be seen in the coastal populations, seems to suggest that the population bottleneck occurred in an east to west direction, followed by differentiation between the north and south in the coastal region. We therefore see strong evidence of Portères’ domestication theory in our population structure analysis.

The fact that *O. glaberrima* does not form a monophyletic clade, however, calls into question the assumption that it speciated through a discrete domestication event. A possible explanation could be the rewilding of ancient *O. glaberrima* landraces, which have ‘gone feral’ and are now classified as the wild species. This would explain the shorter branch lengths of some of these closely related *O. barthii* accessions. Alternatively, one might be tempted to conclude that the paraphyletic nature of this group disproves its taxonomic status as a separate species, and that *O. glaberrima* and *O. barthii* are genetically indistinguishable. Indeed, *O. glaberrima* and *O. barthii* diverged so recently that hybridisation is still considered possible – if difficult – and that admixture between *O. barthii* and *O. glaberrima* even now should not be excluded. In fact, ‘weedy’ rice, which is a genetic mix between the wild and cultivated species, can result from interspecific crosses and has been observed in the case of African rice in both Mali and Cameroon [11].

Another explanation for the observed phylogenetic patterns might be that the time that has passed since domestication (roughly 3000 years ago) has not been sufficient to establish complete lineage sorting. This could mean that *O. glaberrima* still contains a part of the ancestral variation that is observed in *O. barthii*. For this reason, gene trees may not correspond to the overall genomic tree. Indeed, it is a widely observed phenomenon that incomplete lineage causes mixed phylogenetic signals [29].

However, the recurrent pattern of gene haplotypes that are restricted to OG-II accessions from the Guinea Highlands suggests that domestication may have followed a different path in this area, possibly through local introgression from wild rice. This could have led to the observed genetic differences in various parts of the genome. Specifically, the ancestral character of a functionally important SNP in *Sh4* has been confirmed independently and proposed to “support the deep and separate roots of domestication practices in the west versus the eastern cultivation range” [30]. The existence of another locus that explains the loss of shattering in these accessions points to the independent selection of multiple variants affecting the same trait in different sub-populations of *O. glaberrima* and is consistent with parallel adaptation.

### Origins of domestication

The results discussed so far thus shed new light on an old controversy concerning the process of plant domestication in Africa, in which two models have traditionally been competing: the rapid transition model proposed by [2] and the non-centric protracted transition model proposed by [8]. Whereas the work by Portères promotes the idea of a primary domestication centre followed by secondary centres where later improvement occurred, Harlan’s model implies diffuse domestication over a long period of time, with multiple centres or no centres at all. Previous studies that have employed genomic data, have supported both sides of this controversy: Wang et al. [18] found evidence for centric domestication, thereby supporting Portères’ hypothesis, whereas Meyer et al. [19] found evidence for the protracted transition model advocated by Harlan.

This study has aimed to resolve some of the confusion introduced in this debate. First of all, it is clear that the previous genomic studies address different aspects of the controversy: Wang et al. [18] merely propose that domestication was centric, but make no claims regarding the rapidity, and Meyer et al. [19] do the opposite; they propose that domestication was protracted, but not necessarily non-centric. Although the present study does not deal with the time scale of domestication, a few remarks about the potential centre(s) of origin can be made.

Based on the sampling origins of *O. glaberrima*, the results are strongly suggestive of an early domestication event in the eastern cultivation range, with subsequent genetic differentiation towards the west. The geographic origin of the closest wild relatives of *O. glaberrima*, however, correspond mostly with the Senegambian and Guinean forest regions. If we assume that these relatives are genuinely *O. barthii* and not ‘rewilded’ or intermediate forms, there are two plausible scenarios.

Either African rice was first domesticated along the coast and subsequently migrated east. In that case, the coastal landraces must have undergone substantial differentiation in order to explain their larger genetic distance from *O. barthii*. Alternatively, the geographic separation of the inland populations and their wild relatives can be explained by domestication in the eastern cultivation range and a subsequent range shift of the wild progenitor from the east to the west.

To ascertain which of the two scenarios (eastern centre of origin and westward migration, or western centre of origin and eastward migration) eventually holds out, more knowledge is needed about the extent to which *O. glaberrima* and *O. barthii* migrated in the past and whether the present sampling locations truly reflect historical populations. In addition, while the sampling locations of the analysed *O. glaberrima* accessions are quite precise, for most *O. barthii* only the country was known and detailed coordinates were not available. This not only prevented accurate knowledge of the collection sites of these accessions, but also the detection of isolation by distance.

Despite these limitations, the marked population structure observed in *O. glaberrima* points to geographic differentiation during or following domestication. The presence of separate OG-II haplotypes in multiple domestication genes might mean that the segregating landraces acquired domestication traits independently of the majority of *O. glaberrima*. Regardless of whether this happened at the onset of domestication or during a secondary wave, the phylogenetic clustering of these haplotypes with different *O. barthii* accessions provides evidence for a separate genetic origin that may have been caused by introgression or domestication from this otherwise seemingly unrelated wild population.

## Conclusions

In light of the findings presented, we can now with some confidence assert that the centric hypothesis of African rice domestication is incorrect, or at least has some serious shortcomings. The diversity analyses unequivocally demonstrate that *O. glaberrima* underwent an extreme bottleneck. To the best of our knowledge, this bottleneck was most probably associated with domestication, thus supporting the rapid transition model. Although there are some indications of positive selection in terms of an excess of high frequency derived alleles, conclusive evidence of hard selective sweeps – especially in relation to known domestication traits – has been elusive. Whether this stems from methodological issues or from the population structure observed in *O. glaberrima* can only be demonstrated with improved knowledge of the demographic history of *O. glaberrima* and additional modelling.

Although it has been shown that cultivated African rice is much less genetically diverse than wild African rice, the ADMIXTURE analysis shows that *O. glaberrima* is all but homogeneous, even compared to *O. barthii*. The subdivision of the species in coastal and inland populations is suggestive of geographic structure, as is the differentiation along a north-south gradient on the coast. Contrary to expectation, moderate isolation by distance was observed in three out of five genetic sub-populations. Due to data limitations, isolation by distance could not be assessed in *O. barthii*.

Phylogenetic results confirm the clustering of *O. glaberrima* within *O. barthii* but shed new light on the relationships between sub-populations of the wild and domesticated species. *O. glaberrima* indeed shares characteristics with a subset of *O. barthii* individuals; however, the majority of the coastal accessions form a monophyletic clade that does not contain any wild relatives. This pattern breaks down when considering phylogenies at the level of individual genes; there we see that some landraces are far removed from the majority of *O. glaberrima* and cluster with a different *O. barthii* sub-population instead. These separate haplotypes demonstrate the divergent evolutionary trajectories among distant sub-populations, most notably the accessions from the Guinean forest region and the Middle and Lower Niger basin.

Whereas this study provides compelling evidence for the origin of African rice in the eastern cultivation range and its diversification along the Atlantic coast of West Africa, the overarching hypothesis that *O. glaberrima* was domesticated in a single and discrete event has to be rejected. The observed population structure is partially consistent with Portères proposed primary and secondary domestication centres. However, evidence of persisting ancestral variation and multiple gene haplotypes among different sub-populations of *O. glaberrima* suggests that important functional traits may have arisen out of parallel evolution or local adaption, rather than single selective sweeps. This is corroborated by the effect of geographic distance on genetic relatedness and by experimental evidence confirming the phenotypic consequences of spatially restricted genetic variation [19,23,30]. Hence, it can be concluded that the centric, rapid transition model of domestication does not tell the whole story of the evolution of *O. glaberrima*. The protracted transition model with multiple domestication centres, or alternatively a polycentric view, might offer a valuable alternative perspective on the observed geographic distribution of genetic variation found in African rice.

### Future directions

Future research into the origins of African rice should investigate the possibility that the closest wild relatives of *O. glaberrima* are in fact hybrids or rewilded ancient landraces. A closer examination of the genetic diversity and signatures of selection of the hypothetical ancestor population in comparison to other *O. barthii* and *O. glaberrima* accessions might elucidate whether they are more similar to the domesticated or to the wild species. These genetic analyses will have to be balanced with suitable morphological evidence. Uncertainty in species delimitation could be further examined through phylogenetic networks and introgression analyses. This will aid our understanding of the precise evolutionary relationships between *O. glaberrima* and *O. barthii*.

In addition, focus should be given to more even sampling across the geographic range of both species, especially in the eastern range for *O. glaberrima* and in the western range for *O. barthii*, where collections of these species are presently scarce. Larger sample sizes will also increase the sensitivity of genome-wide association studies, which enable the identification of SNPs that are associated with traits of interest. While genome-wide data have already been used to explore the mutations associated with drought tolerance [19], these data have not yet been mined for other signs of ecological adaptation.

Lastly, more functional analyses are needed to improve the annotation of the African rice genome, which is still lacking in many ways in comparison to the Asian rice genome. This will help to predict the phenotypic consequences of gene haplotypes and in linking phylogenetic patterns to the evolution of functionally significant traits. The complementation of computational studies with experimental data will be indispensable in the future – not just for understanding the broad patterns of evolution and domestication of African rice, but also to provide insights into the emergence of local adaptive traits connected with the diversification of this crop in its different geographic contexts.

## Materials and Methods

### Whole genome alignment and variant discovery

This study used publicly available whole genome data of 111 *O. glaberrima* and 94 *O. barthii* accessions. A list of all used accessions and their metadata can be found in S1 Table. Variants were called relative to the *O. glaberrima* (AGI1.1) reference genome and subsequently filtered to remove false positives. The *O. glaberrima* reference genome [18] was retrieved from Ensembl Genomes (release 33). Variant discovery was performed following the Genome Analysis Tool Kit (GATK) Best Practices [31]. Untrimmed reads were mapped to the reference genome using the BWA-MEM algorithm [32] of the Burrows-Wheeler Aligner (v0.7.13). Duplicate reads were flagged with Picard (v1.129) MarkDuplicates [33]. Local realignment was performed around indels with GATK (v3.6.0) RealignerTargetCreator and IndelRealigner [34]. The resulting BAM files were indexed and validated with Picard (v1.129). Individual genotypes were called using GATK (v3.6.0) HaplotypeCaller on reads with a minimum mapping quality score of 30. GVCFs were combined into a single VCF with GATK (v3.6.0) GenotypeGVCFs. Only biallelic SNPs were retained for analysis.

### Quality filtering

Hard filters were applied to the raw SNPs by removing SNPs falling outside the quality thresholds of several common annotations: DP, QD, MQ, MQRankSum, ReadPosRankSum and FS (for an explanation, see S11 Table). In addition, we used the no-call rate divided by the number of samples as a measure of missing data. We did not filter for heterozygous sites, because both species are primarily inbreeding and therefore exhibit low levels of heterozygosity. Filter thresholds were determined based on their effect on the Transition:Transversion ratio (Ts:Tv). Although the true Ts:Tv ratio is unknown and varies along the genome, it is known that functional constraints generally favour transitions over transversions [35], leading to ‘transition bias’. SNPs with a higher Ts:Tv are thus likely to be enriched for true SNPs, while SNPs with a lower Ts:Tv will contain more false positives. SNPs were binned along the range of a given annotation. For each interval, SNP count and Ts:Tv were calculated and plotted in R (v3.3.2) [36]. Intervals were removed in order to increase Ts:Tv while retaining a reasonable number of SNPs. Based on previous studies, call sets between 2 and 4 million SNPs were deemed reasonable.

Filtering criteria with different levels of strictness were applied (S12 Table). This resulted in two call sets. The first call set contained a full set of SNPs considered to adhere to a minimum standard of quality. The second call set contained fewer SNPs that adhere to a higher standard of quality. Both call sets were used; a choice between the two was made depending on the amount and quality of SNPs needed for each analysis. Relative diversity estimates, selection scans and detection of population structure require fewer SNPs; for those analysis, the reduced call set was used. In order to obtain pairwise genomic distances and differentiate between gene haplotypes, a higher density of SNPs was desired; for these analyses, the complete call set was used, so as to maximise the number of segregating sites. The effect of filtering on Ts:Tv ratio was quantified with VCFTools (v0.1.14) [37]. Filter classes and their thresholds can be found in S13 Fig.

### Variant statistics

Mean depth of coverage, fraction of missing data and mean variant quality per SNP were calculated in 100 kb sliding windows along the entire genome in order to assess the distribution and quality of SNPs. These statistics were computed using VCFtools (v0.1.14) for both call sets. Large problematic regions were not detected. In order to make an informed decision as to which version of call set to use and whether or not to adjust the filtering parameters, we calculated additional statistics. Following the method of [22], both call sets were compared with respect to their patterns of nucleotide diversity (π) and genetic differentiation (F_ST_). The fixation index (F_ST_) between *O. glaberrima* and *O. barthii* is based on the implementation of [38] and calculated as: 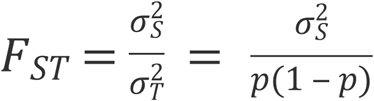, where *p* is the allele frequency in the total population, *σ*^*2*^_*T*_ is the variance in allele frequency in the total population, and *σ*^*2*^_*S*_ is the variance in allele frequency between the two sub-populations. Relative nucleotide diversity was calculated as the ratio of π in *O. glaberrima* to π in *O. barthii*, where π defined as:

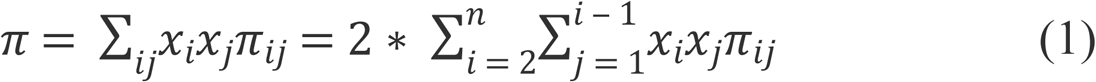

Here π_*ij*_ is the number of differences per site between sequences *i* and *j*, *x*_*i*_ is the frequency of sequence *i, x*_*j*_ is the frequency of sequence *j* and *n* is the total number of sequences in the data set. The results were deemed sufficiently comparable to proceed with both call sets (S14 Fig). In addition, site depth, call rate and mean heterozygosity per individual were calculated for all accessions using VCFtools (v0.1.14). An overview of these statistics and the number and types of SNPs in the two sets can be found in S15 Table.

In order to estimate the genetic diversity in *O. barthii* and *O. glaberrima* separately, variants were split into two populations based on species identification. Monomorphic sites were removed. A total of 2,580,362 and 1,419,601 SNPs were used to calculate SNP density, π, and Tajima’s D in *O. barthii* and *O. glaberrima*, respectively, where Tajima’s D is defined according to [39] and measures the difference between two estimators of ϴ (the scaled mutation rate), namely the average number of differences between two sequences (π) as per Equation (1) and the expected number of segregating sites between two sequences under neutral theory according to Watterson’s estimator 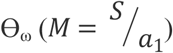, where *S* is the total number of segregating sites in the population, 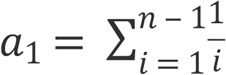, and *i* is the *i*^th^ sequence in a total of *n* sequences. When population size is constant and there is no selection on the genome (so-called neutral conditions), the two estimators should equal each other and Tajima’s D equals 0. These statistics were computed in 100 kb regions with VCFtools (v0.1.14). Genome-wide average statistics were compared using the Kruskal-Wallis test. Minor allele frequencies were calculated using VCFtools (v0.1.14) in combination with a custom R script and plotted in R (v.3.3.2).

### Derived allele frequency spectrum

*Oryza* is a large genus and many reference genomes of related species are available. Three potential outgroups were identified to polarise SNPs, in order of decreasing genetic distance to *O. glaberrima*: *Oryza punctata, Oryza meridionalis,* and *Oryza longistaminata* A. Chev. & Roehr. (S16 Table). These outgroups were determined based on known divergence times and genomic distances to *O. sativa* ssp. *japonica* which, as a sister taxon, is supposed to be equidistant to the outgroup as compared to *O. glaberrima.* For the genetic distances reported to the candidate species, saturation analyses show a good fit between the corrected pairwise divergence and the uncorrected P-distance [40]. For this reason, homoplasy was considered unlikely and correction for multiple substitutions was not applied. A closely related outgroup offers a better alignment but carries the risk of incomplete lineage sorting and therefore incorrectly assigned ancestral alleles. A distantly related outgroup circumvents this problem but is more difficult to align, and hence will cause larger loss of data. *Oryza longistaminata* was rejected because of its low genomic divergence from *O. glaberrima* (∼2%). *Oryza punctata* was rejected because of its high genomic divergence (>5%) and associated data loss. Hence, we chose *O. meridionalis* as an outgroup.

SNPs were polarised with reference to the *O. meridionalis* (v1.3) genome [41] with a custom R script. The *O. meridionalis x O. glaberrima* multiple alignment was retrieved from Ensembl Genomes (release 33) and parsed with mafTools [42]. For each biallelic SNP, the corresponding position and five flanking bases were extracted from the alignment using a custom perl script. Positions that did not map to the outgroup, positions with gaps within 5 bp of the SNP, and SNPs that mapped to multiple regions of the *O. meridionalis* genome were discarded. A total of 3,923,601 variants were screened, of which 2,332,467 either did not align to the outgroup species or did not pass the alignment quality filter. The derived allele frequency spectrum was calculated for all synonymous and noncoding variants of the remaining 1,591,134 SNPs.

Synonymous and non-coding SNPs were extracted using SnpSift (v4.0) [43]. For these SNPs, the derived allele frequency spectra and cumulative densities were plotted using the R (v3.3.2). Because of the low genomic divergence (<5%), homoplasy was considered unlikely and correction for multiple substitutions was not applied. The expected site frequency spectrum under a neutral model of evolution was calculated using the estimation of the population scaled mutation rate [39]. Deviation from neutrality of the observed site frequency spectra of the two populations was compared using a two-sample Kolmogorov-Smirnov test.

### Selection scans

Several CLR tests that are widely used for detecting ‘hard’ sweeps are available as open source software, including OmegaPlus [44], SweeD [45] and SweepFinder [46]. These CLR methods are superior to more common neutrality tests such as Tajima’s D, because they measure deviations of the site frequency spectrum (SFS) against the genomic ‘background’ SFS. Since the background has been partly shaped by demographic history, these scans thereby each to some extent control for the confounding effect of past fluctuations in population size. In comparative analyses of a number of these CLR methods using simulated data, SweeD and OmegaPlus were shown to outperform other tests [47]. While SweeD is capable of taking into account the polarisation of alleles in the so-called ‘unfolded’ SFS, this has the disadvantage that limiting the analyses to only unfolded SNPs causes a significant loss of data. OmegaPlus has no feature to distinguish between ancestral and derived alleles, but has the added advantage of explicitly taking into account patterns of Linkage Disequilibrium (LD). Extensive LD is observed in the *O. glaberrima* genome, with *r*^*2*^ reaching half its maximum value at a distance of 175 kb and approaching baseline at 300 kb [19]. For these reasons, OmegaPlus was chosen as the preferred method.

The ω test statistic is calculated as:

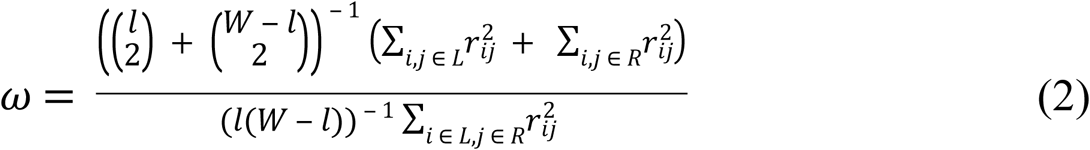

Here *W* is the number of segregating sites, divided into two groups: one from the first to the *l*^th^ polymorphic site on the left and the other from the (*l* + 1)^th^ to the last polymorphic site on the right. *L* and *R* represent the left and right set of polymorphic sites, respectively, and *r*^*2*^ _*ij*_ is *r*^*2*^, a common measure of LD [48], between the *i*^th^ and the *j*^th^ site. The value of *l* that maximises ω defines the test statistic. (Note: LD is defined by Hill and Robertson [48] as *r*^*2*^ *= D*/*p*_1_*p*_2_*q*_1_ *q*_2_, where *p*_*1*_ and *p*_*2*_ are the allele frequencies of SNP1, *q*_*1*_ and *q*_*2*_ are the allele frequencies of SNP2, and *D* measures the absolute difference between the observed and the expected haplotype frequencies *p*_*1*_*q*_*1*_, *p*_*2*_*q*_*1*_, *p*_*1*_*q*_*2*_ and *p*_*2*_*q*_*2*_ respectively.)

Values of ω were log-transformed, prior to creating Manhattan plots in R (v3.3.2) using the ‘qqman’ package [49]. Windows containing domestication genes with previous evidence for positive selection were highlighted. Positions within the top 0.5% values were considered candidate regions. To verify whether these candidate regions show other characteristic signatures of selection, the ω-statistic was plotted against overlapping 25 kb windows of π and Tajima’s D, respectively (S5 Fig). Since common outliers were rare and would require lowering the threshold for OmegaPlus, we did not report any common outliers as candidate regions, but rather chose OmegaPlus as the leading test.

Candidate regions were screened for potential causative mutations by examining related SNP content and genomic features. Variants were annotated using SnpEff (v4.0) [43]. Genomic features were retrieved from the general feature format (GFF) file of the *O. glaberrima* reference genome on Ensembl (release 33). Genomic features containing putative moderate to high impact mutations within close proximity (< 25 kb) of candidate regions were extracted for closer inspection using a custom R script. Candidate sweeps (OmegaPlus outliers) and their associated genes harbouring high impact mutations can be found in S17-S19 Tables.

### Population structure

Population structure was determined with ADMIXTURE (v1.3.0) [50]. In order to minimise the confounding effect of linkage disequilibrium, SNPs with a correlation coefficient of r^2^ > 0.25 were pruned with PLINK (v2.0) in sliding windows of 500 SNPs, with a step size of 50 SNPs [51]. A total of 70,873 SNPs were retained for analysis. ADMIXTURE was subsequently run with varying levels of K and cross-validation to improve model fit. An optimal number of ancestral populations was selected by choosing the level of K with the lowest cross-validation error. This analysis was repeated for *O. glaberrima* and *O. barthii* separately. Cross-validation error estimates of all three analyses can be found in S6 Fig. The resulting ancestry fractions were plotted as stacked bar charts in R (v3.3.2). The *O. glaberrima* population was subsequently divided into five populations based on the ancestral population contributing the highest fraction of genetic variation.

The geographic distribution of these populations in West Africa was visualised by plotting the coordinates of all West African accessions using the R packages ‘rworldmap’ and ‘raster’ [52,53]. The top 20 principal components were calculated for both species together and for both species separately using PLINK (v1.9). The correlation of the top two principal components with longitude and latitude was calculated for all accessions collected from West Africa with known coordinates. The significance of the correlation coefficient was assessed by conducting a two-sided t-test, under the null hypothesis that the top two principal components are not significantly correlated with either latitude or longitude.

### Geographic differentiation

Based on the results by Meyer et al. [19], four geographic sub-populations were identified, separating the arid and tropical populations by the 11° N cline and coastal and inland populations by the 6° W cline, respectively. Allelic differentiation between these populations was measured using Weir and Cockerham’s [38] definition of F_ST_, as implemented in VCFtools (v0.1.14). F_ST_ between *O. glaberrima* and *O. barthii* was included as a baseline. To account for uneven sample sizes of these populations, an equal number of individuals (n=15) was selected for each of the five sub-populations as identified with ADMIXTURE. To minimise the effect of missing data, an equal number of individuals (n=15) that were sequenced at high coverage were selected from the *O. barthii* population. Pairwise F_ST_ and π were calculated between all six groups to obtain a more balanced estimate of allelic differentiation, taking uneven sampling and sequencing depth into account.

Isolation by distance (IBD) was assessed by comparing the relatedness of individuals with the geographic distance separating their sites of collection. Kinship coefficients were estimated based on 1,419,601 SNPs, using the KING robust relationship inference method [54] as implemented in PLINK (v2.0) [51]. Pairwise geographic distances were calculated in R (v3.3.2) using the package ‘geosphere’ [55], as the shortest distance between two points according to the Haversine function, assuming a spherical Earth with a radius of 6,378 km. Box and whisker plots of the resulting distances and kinships were grouped per genetic cluster and visualised in R (v3.3.2). The relation between kinship and geographic distance was quantified by fitting a linear model to the data points. To reflect the geographic range of the majority of each genetic population, outliers were omitted. A pair was considered an outlier when the distance separating them fell outside the interquartile range (IQR) by more than 1.5*IQR. On average, outliers comprised less than 4% of the data. Linear models and correlation coefficients were estimated in R (v3.3.2). Kinship by distance graphs for each population separately and all populations combined can be found in S7 Fig.

### Phylogenetic analyses

To confirm the clustering of *O. glaberrima* within *O. barthii*, a whole genome phylogenetic tree was constructed based on 3,923,601 genome-wide SNPs. To avoid distortion of branch lengths, no outgroup was used. Pairwise genomic distances between all accessions were calculated using a custom perl script, implementing the method described in S3.2 of [56]. The divergence between two genomes X and Y was calculated as:

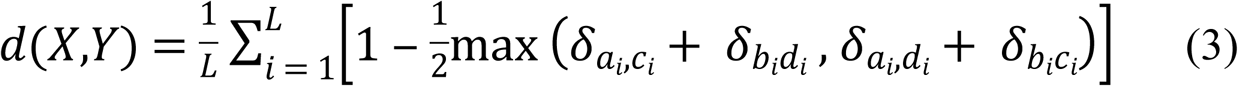

Here *a*_*i*_*b*_*i*_ is the genotype at position *i* in *X*, and *c*_*i*_*d*_*i*_ is the genotype at position *i* in *Y*. The resulting distance matrix was used to construct an unrooted neighbour joining (NJ) tree, using the BioNJ algorithm of FastME (v2.0) with subtree pruning and regrafting (SPR) [57]. Trees were pruned and annotated in Interactive Tree Of Life (iTOL v3) [58].

Gene trees were constructed of salient domestication genes that have been proposed to play a role in rice evolution [18,22]. These genes were identified according to criteria published in Meyer & Purugganan [59]. The selected domestication genes and their proposed functions are listed in S8 Table. Because the African rice genome is poorly annotated, their putative location in the *O. glaberrima* genome was determined using the *O. sativa* reference genome as a guide. *O. sativa* protein sequences were retrieved from the Rice Annotation Project Database (RAP-DB) and compared against the Ensembl Genomes *O. glaberrima* reference protein FASTA using BLAST+ (v2.6) [60,61]. Genes were considered homologous when protein sequence similarity was higher than or equal to 95%. Based on this criterion, 19 out of 25 genes could be used for analysis. Gene coordinates were retrieved from the GFF file of the *O. glaberrima* reference genome on Ensembl (release 33). Gene structures of these genes were retrieved from Ensembl Plants (release 37).

To ensure sufficient phylogenetic signal, gene intervals were extended with 5 kb flanking regions on either side using bedtools slop (v2.26.0) [62]. Larger intervals were not used, to minimise the influence of LD decay. The SNPs within the selected genomic coordinates were phased with PHASE (v2.1.1) [63,64], using the general model for recombination rate variation as found in Li & Stephens [65]. Output was converted to FASTA format with a custom perl script. The resulting multiple alignments each contained 412 nucleotide sequences. Evolutionary histories were inferred using the NJ algorithm [66] as implemented in MEGA7 [67]. Both coding and noncoding positions were used. Evolutionary distances were calculated using the p-distance method of Nei & Kumar [68]. Ambiguous positions were removed for each sequence pair. All trees were annotated in Interactive Tree Of Life (iTOL v3) [58]. The five most common haplotypes for each gene were identified using the PHASE output. Putative effects of segregating SNPs were predicted using SnpEff (v4.0). Tree and haplotype statistics of all genes are summarised in S9 Table.

## Acknowledgements

This work was made possible by generous support from the Biosystematics group at Wageningen University and the Purugganan lab at New York University. Sincere thanks to all members of the lab in New York, specifically Michael and Jae, for many useful discussions.

## Supporting information

**S1 Table. List of accessions included in this study S2 Fig. Geographic origin of used accessions**

**S3 Fig. Relative genetic diversity and allele frequencies in domesticated and wild rice**

**S4 Fig. Log-transformed CLR test statistic (ω)**

**S5 Fig. Correspondence between ω and other neutrality tests**

**S6 Fig. Cross-validation (CV) error estimates of ADMIXTURE, with varying levels of K**

**S7 Fig. Isolation by distance in the five genetic clusters of *O. glaberrima***

**S8 Table. Genes selected for phylogenetic analysis**

**S9 Table. Phasing and phylogenetic output of domestication genes**

**S10 Fig. Separation of OG-IV haplotype in multiple domestication genes**

**S11 Table. Description of quality control filters**

**S12 Table. Quality control filter thresholds**

**S13 Fig. SNP count and Ts:Tv ratio grouped by filter class**

**S14 Fig. Effect of filtering thresholds on nucleotide diversity and fixation index**

**S15 Table. Summary of variant calls**

**S16 Table. Potential outgroup species**

**S17 Table. Candidate selective sweeps unique to *O. glaberrima***

**S18 Table. Genes associated with candidate sweeps**

**S19 Table. High impact mutations associated with candidate sweeps**

## References

1. Molina J, Sikora M, Garud N, Flowers JM, Rubinstein S, Reynolds A, et al. Molecular evidence for a single evolutionary origin of domesticated rice. Proc Natl Acad Sci U S A. 2011 May 17;108(20):8351–6.

2. Porteres R. Berceaux Agricoles Primaires Sur le Continent Africain. J Afr Hist. 1962;3(2):195–2010.

3. Linares OF. African rice (Oryza glaberrima): history and future potential. Proc Natl Acad Sci U S A. 2002 Dec 10;99(25):16360–5.

4. Van Andel T. African Rice (Oryza glaberrima Steud.): Lost Crop of the Enslaved Africans Discovered in Suriname. Econ Bot. 2010 Mar;64(1):1–10.

5. Van Andel TR, Van der Velden A, Reijers M. The ‘Botanical Gardens of the Dispossessed’ revisited: richness and significance of Old World crops grown by Suriname Maroons. Genet Resour Crop Evol. 2016 Apr 9;63(4):695–710.

6. Mohanty S. IRRI - Trends in global rice consumption [Internet]. 2013 [cited 2017 May 30]. Available from: http://irri.org/rice-today/trends-in-global-rice-consumption

7. Schmidhuber J, Tubiello FN. Global food security under climate change. Proc Natl Acad Sci U S A. 2007 Dec 11;104(50):19703–8.

8. Harlan JR, De Wet JMJ, Stemler A. Plant Domestication and Indigenous African Agriculture. In: Origins of African Plant Domestication. De Gruyter Mouton; 1976. p. 3–19.

9. Clark JD. The Problem of Neolithic culture in sub-Saharan Africa. In: Bishop WW, Clark JD, editors. Background to Evolution in Africa. Chicago, IL: Chicago University Press; 1967. p. 601–27.

10. Shaw T. Early crops in Africa: a review of the evidence. In: Harlan JR, De Wet JMJ, Stemler ABL, editors. Origins of African Plant Domestication. The Hague & Paris: Mouton; 1976. p. 107–53.

11. Orjuela J, Sabot F, Chéron S, Vigouroux Y, Adam H, Chrestin H, et al. An extensive analysis of the African rice genetic diversity through a global genotyping. Theor Appl Genet. 2014 Oct;127(10):2211–23.

12. Gross BL, Zhao Z. Archaeological and genetic insights into the origins of domesticated rice. Proc Natl Acad Sci U S A. 2014 Apr 29;111(17):6190–7.

13. Allaby RG. Barley domestication: the end of a central dogma? Genome Biol. 2015 Dec 26;16(1):176.

14. Choi JY, Platts AE, Fuller DQ, Hsing Y-I, Wing RA, Purugganan MD. The rice paradox: Multiple origins but single domestication in Asian rice. Mol Biol Evol. 2017 Jan 12;34(4):msx049.

15. Li Z-M, Zheng X-M, Ge S. Genetic diversity and domestication history of African rice (Oryza glaberrima) as inferred from multiple gene sequences. Theor Appl Genet. 2011 Jun 12;123(1):21–31.

16. Semon M, Nielsen R, Jones MP, McCouch SR. The population structure of African cultivated rice oryza glaberrima (Steud.): evidence for elevated levels of linkage disequilibrium caused by admixture with O. sativa and ecological adaptation. Genetics. 2005 Mar 1;169(3):1639–47.

17. Nabholz B, Sarah G, Sabot F, Ruiz M, Adam H, Nidelet S, et al. Transcriptome population genomics reveals severe bottleneck and domestication cost in the African rice (Oryza glaberrima). Mol Ecol. 2014 May;23(9):2210–27.

18. Wang M, Yu Y, Haberer G, Marri PR, Fan C, Goicoechea JL, et al. The genome sequence of African rice (Oryza glaberrima) and evidence for independent domestication. Nat Genet. 2014 Sep 27;46(9):982–8.

19. Meyer RS, Choi JY, Sanches M, Plessis A, Flowers JM, Amas J, et al. Domestication history and geographical adaptation inferred from a SNP map of African rice. Nat Genet. 2016 Aug 8;48(9):1083–8.

20. Hartl DL, Clark GC. Principles of Population Genetics. Sunderland: Sinauer Associates; 1997.

21. Ishida Y. Sewall Wright and Gustave Malécot on Isolation by Distance. Philos Sci. 2009;76(5):784–96.

22. Li L-F, Li Y-L, Jia Y, Caicedo AL, Olsen KM. Signatures of adaptation in the weedy rice genome. Nat Genet. 2017 Apr 3;49(5):811–4.

23. Lv S, Wu W, Wang M, Meyer RS, Ndjiondjop M-N, Tan L, et al. Genetic control of seed shattering during African rice domestication. Nat Plants. 2018;4:331–7.

24. Ohnishi T, Sugahara S, Yamada T, Kikuchi K, Yoshiba Y, Hirano H-Y, et al. OsNAC6, a member of the NAC gene family, is induced by various stresses in rice. Genes Genet Syst. 2005 Apr;80(2):135–9.

25. Nielsen R. Molecular signatures of natural selection. Annu Rev Genet. 2005;39:197–218.

26. Ndjiondjop M-N, Semagn K, Gouda AC, Kpeki SB, Dro Tia D, Sow M, et al. Genetic Variation and Population Structure of Oryza glaberrima and Development of a Mini-Core Collection Using DArTseq. Front Plant Sci. 2017 Oct 17;8:1748.

27. Pfaffelhuber P, Lehnert A, Stephan W, Parsch J. Linkage Disequilibrium Under Genetic Hitchhiking in Finite Populations. Genetics. 2008 May 1;179(1):527–37.

28. Ralph P, Coop G. Parallel adaptation: one or many waves of advance of an advantageous allele? Genetics. 2010 Oct 1;186(2):647–68.

29. Degnan JH, Rosenberg NA. Gene tree discordance, phylogenetic inference and the multispecies coalescent. Trends Ecol Evol. 2009;24(6):332–40.

30. Wu W, Liu X, Wang M, Meyer RS, Luo X, Ndjiondjop M-N, et al. A single-nucleotide polymorphism causes smaller grain size and loss of seed shattering during African rice domestication. Nat Plants. 2017 May 8;3(6):17064.

31. DePristo MA, Banks E, Poplin R, Garimella K V, Maguire JR, Hartl C, et al. A framework for variation discovery and genotyping using next-generation DNA sequencing data. Nat Genet. 2011 May 10;43(5):491–8.

32. Li H, Durbin R. Fast and accurate long-read alignment with Burrows-Wheeler transform. Bioinformatics. 2010;26(5):589–95.

33. Broad Institute. Picard Tools [Internet]. [cited 2017 Aug 7]. Available from: http://broadinstitute.github.io/picard/

34. McKenna A, Hanna M, Banks E, Sivachenko A, Cibulskis K, Kernytsky A, et al. The Genome Analysis Toolkit: a MapReduce framework for analyzing next-generation DNA sequencing data. Genome Res. 2010 Sep 1;20(9):1297–303.

35. Li WH, Wu CI, Luo CC. A new method for estimating synonymous and nonsynonymous rates of nucleotide substitution considering the relative likelihood of nucleotide and codon changes. Mol Biol Evol. 1985 Mar;2(2):150–74.

36. R Core Team. R: A language and environment for statistical computing. Vienna, Austria: R Foundation for Statistical Computing; 2013.

37. Danecek P, Auton A, Abecasis G, Albers CA, Banks E, DePristo MA, et al. The variant call format and VCFtools. Bioinformatics. 2011;27(15):2156–8.

38. Weir BS, Cockerham CC. Estimating F-Statistics for the Analysis of Population Structure. Evolution (N Y). 1984;38(6):1358–70.

39. Tajima F. Statistical method for testing the neutral mutation hypothesis by DNA polymorphism. Genetics. 1989 Nov;123(3):585–95.

40. Zhu T, Xu P-Z, Liu J-P, Peng S, Mo X-C, Gao L-Z. Phylogenetic relationships and genome divergence among the AA-genome species of the genus Oryza as revealed by 53 nuclear genes and 16 intergenic regions. Mol Phylogenet Evol. 2014 Jan;70:348–61.

41. Jacquemin J, Bhatia D, Singh K, Wing RA. The International Oryza Map Alignment Project: development of a genus-wide comparative genomics platform to help solve the 9 billion-people question. Curr Opin Plant Biol. 2013 May 1;16(2):147–56.

42. Earl D, Paten B, Diekhans M. Alignathon: a competitive assessment of whole-genome alignment methods. Genome Res. 2014;24(12):2077–89.

43. Cingolani P, Patel VM, Coon M, Nguyen T, Land SJ, Ruden DM, et al. Using Drosophila melanogaster as a model for genotoxic chemical mutational studies with a new program, SnpSift. Front Genet. 2012;3(35).

44. Kim Y, Nielsen R. Linkage Disequilibrium as a Signature of Selective Sweeps. Genetics. 2004;167(3).

45. Pavlidis P, Zivkovic D, Stamatakis A, Alachiotis N. SweeD: Likelihood-Based Detection of Selective Sweeps in Thousands of Genomes. Mol Biol Evol. 2013 Jun 18;30(9):2224–34.

46. Nielsen R, Williamson S, Kim Y, Hubisz MJ, Clark AG, Bustamante C. Genomic scans for selective sweeps using SNP data. Genome Res. 2005 Nov;15(11):1566–75.

47. Pavlidis P, Alachiotis N. A survey of methods and tools to detect recent and strong positive selection. J Biol Res. 2017 Dec 8;24(1):7.

48. Hill WG, Robertson A. Linkage Disequilibrium in Finite Populations. Theor Appl Genet. 1968;38:226–23.

49. Turner SD. qqman: an R package for visualizing GWAS results using Q-Q and manhattan plots. biorXiv.

50. Alexander DH, Novembre J, Lange K. Fast model-based estimation of ancestry in unrelated individuals. Genome Res. 2009;19:1655–1664.

51. Purcell S, Neale B, Todd-Brown K, Thomas L, Ferreira MAR, Bender D, et al. PLINK: a tool set for whole-genome association and population-based linkage analyses. Am J Hum Genet. 2007 Sep;81(3):559–75.

52. Hijmans RJ, Van Etten J. raster: Geographic analysis and modeling with raster data. 2012.

53. South A. rworldmap: A New R package for Mapping Global Data. R J. 2011;3(1):35–43.

54. Manichaikul A, Mychaleckyj JC, Rich SS, Daly K, Sale M, Chen W-M. Robust relationship inference in genome-wide association studies. Bioinformatics. 2010 Nov 15;26(22):2867–73.

55. Hijmans RJ. geosphere: Spherical Trigonometry. R package version 1.5-5. 2016.

56. Gronau I, Hubisz MJ, Gulko B, Danko CG, Siepel A. Bayesian inference of ancient human demography from individual genome sequences. Nat Genet. 2011 Sep 18;43(10):1031–4.

57. Lefort V, Desper R, Gascuel O, M A, W H, O G. FastME 2.0: A Comprehensive, Accurate, and Fast Distance-Based Phylogeny Inference Program: Table 1. Mol Biol Evol. 2015 Oct 1;32(10):2798–800.

58. Letunic I, Bork P. Interactive tree of life (iTOL) v3: an online tool for the display and annotation of phylogenetic and other trees. Nucleic Acids Res. 2016;

59. Meyer RS, Purugganan MD. Evolution of crop species: genetics of domestication and diversification. Nat Rev Genet. 2013 Nov 18;14(12):840–52.

60. Camacho C, Coulouris G, Avagyan V, Ma N, Papadopoulos J, Bealer K, et al. BLAST+: architecture and applications. BMC Bioinformatics. 2009;(10):421.

61. Sakai H, Lee SS, Tanaka T, Numa H, Kim J, Kawahara Y, et al. Rice Annotation Project Database (RAP-DB): An Integrative and Interactive Database for Rice Genomics. Plant Cell Physiol. 2013 Feb;54(2):e6–e6.

62. Quinlan AR, Hall IM. BEDTools: a flexible suite of utilities for comparing genomic features. Bioinformatics. 2010;26(6):841–84210.

63. Stephens M, Smith NJ, Donnelly P. A New Statistical Method for Haplotype Reconstruction from Population Data. Am J Hum Genet. 2001;68:978–89.

64. Stephens M, Donnelly P. A Comparison of Bayesian Methods for Haplotype Reconstruction from Population Genotype Data. Am J Hum Genet. 2003;73:1162–9.

65. Li N, Stephens M. Modeling Linkage Disequilibrium and Identifying Recombination Hotspots Using Single-Nucleotide Polymorphism Data. Genetics. 2003;165:2213–33.

66. Saitou N, Nei M. The neighbor-joining method: a new method for reconstructing phylogenetic trees. Mol Biol Evol. 1987 Jul;4(4):406–25.

67. Kumar S, Stecher G, Tamura K. MEGA7: Molecular Evolutionary Genetics Analysis version 7.0 for bigger datasets. Mol Biol Evol. 2016;33:1870–4.

68. Nei M, Kumar S. Molecular Evolution and Phylogenetics. New York: Oxford University Press; 2000.

